# Modulation of heteromeric glycine receptor function through high concentration clustering

**DOI:** 10.1101/2024.10.17.618879

**Authors:** Hailong Yu, Weiwei Wang

## Abstract

Ion channels are targeted by many drugs for treating neurological, musculoskeletal, renal and other diseases. These drugs bind to and alter the function of individual channels to achieve desired therapeutic effects. However, many ion channels function in high concentration clusters in their native environment. It is unclear if and how clustering modulates ion channel function. Human heteromeric glycine receptors (GlyRs) are the major inhibitory neurotransmitter receptors in the spinal cord and are active targets for developing chronic pain medications. We show that the α2β heteromeric GlyR assembles with the master postsynaptic scaffolding gephyrin (GPHN) into micron-sized clustered at the plasma membrane after heterologous expression. The inhibitory trans-synaptic adhesion protein neuroligin-2 (NL2) further increases both the cluster sizes and GlyR concentration. The apparent glycine affinity increases monotonically as a function of GlyR concentration but not with cluster size. We also show that ligand re-binding to adjacent GlyRs alters kinetics but not chemical equilibrium. A positively charged N-terminus sequence of the GlyR β subunit was further identified essential for glycine affinity modulation through clustering. Taken together, we propose a mechanism where clustering enhances local electrostatic potential, which in turn concentrates ions and ligands, modulating the function of GlyR. This mechanism is likely universal across ion channel clusters found ubiquitously in biology and provides new perspectives in possible pharmaceutical development.

## Introduction

Ion channels underlie both vital bodily functions such as heartbeat, muscle control and hormone secretion, as well as advanced cognitive abilities including emotion, learning, and memory^1–4^. Dysregulation and/or genetic mutations of ion channels lead to a myriad of diseases and conditions including pain, cardiac arrythmia, muscle dysfunction, renal cysts, cancer, epilepsy, dementia, and other problems^5–7^. Modulation of ion channel function clearly has strong physiological and therapeutical implications, making them the second largest drug targets following GPCRs^1,4,8^.

Although much has been learned of the working mechanism and ligand regulation of individual ion channels, many channels do not function in isolation, but form high concentration clusters in native environments. Examples of clustered channels include ionotropic neurotransmitter receptors at synapses^2,9–11^, acetylcholine receptors at neuromuscular junctions^12,13^, voltage gated Na^+^ and K^+^ channels at the nodes of Ranvier^14–17^, and Ca^2+^ channels in the plasma/sarcoplasmic/endoplasmic membranes^18–20^. Clustering has been reported to induce cooperative gating through common ligands (usually Ca^2+^), or through presumable direct mechanical coupling for several channels ^21,22^. However, it is unclear if and how clustering of neurotransmitter receptors affect function. In addition, whether there are more general mechanisms of clustering-induced functional modulation is unclear.

Glycine receptors (GlyRs) are the major inhibitory neurotransmitter receptors in the spinal cord, the dysfunction of which causes hyperekplexia^23–26^ and neuropathic inflammatory pain^27–29^. Regulators of GlyRs, including cannabinoids and propofol derivatives, have shown strong potential in pain treatment^29–34^. Recent identification of the molecular composition^35,36^, working mechanisms^37,38^ and drug modulation^39^ of human heteromeric GlyRs has provided basis for understanding how individual channels work. However, in neurons, heteromeric GlyRs are clustered at high concentrations, up to 10, 000 GlyR channels per µm^2^, at post-synaptic membranes through direct interaction with the scaffolding protein gephyrin (GPHN) by the GlyR β subunit^10,40–44^. It is unclear if and how high-concentration clustering modulates GlyR function.

In this work, we recapitulated the clustering of a human heteromeric GlyR, the α2β GlyR, by GPHN in HEK293 cells. In addition, we found that the trans-synaptic adhesion molecule neuroligin-2 (NL2), promoted the clustering in both size and GlyR concentration. Whole-cell electrophysiology recordings showed higher glycine affinity upon clustering. Through mutagenesis of GPHN, and quantitative analysis of total internal reflection fluorescence (TIRF) micrographs of unroofed cells, we show that the increase of glycine affinity is monotonically correlated with higher GlyR concentrations in clusters. We further show that fast re-binding in high concentration clusters slowed ligand interaction kinetics but did not shift equilibrium, while local accumulation of electrostatic charges carried by the GlyR β subunit upon clustering increased glycine affinity. Since charged sequences are commonly found in ion channels, we propose that the enhancement of local electrostatic potential through clustering may be universal mechanism for ion channel function modulation.

## RESULTS

### Clustering of α2β GlyR requires full-length gephyrin

The GPHN E domain (GPHN_E) binds with GlyR at plasma membrane but does not induce clustering. GPHN is composed of the E domain that binds to GlyR, the G domain that forms a trimer, and the semi-flexible C domine that changes compactness upon phosphorylation^45–49^ (Fig. 1A bottom). We generated GlyR with a single eGFP fusion on the β subunit for fluorescence imaging and quantification of fluorescence intensity from one GlyR (Fig. S1A, taking account the low oligomers reported before, see methods)^35,50^. GPHN and GPHN_E are fused to mCherry for detection. Heterologous expression of the α2β GlyR alone led to diffraction-limited fluorescence spots in micrographs of unroofed cell membranes imaged using TIRF (Fig. 1B, see methods for details). Co-expression of GPHN_E and GlyR yielded mostly co-localizing (Fig. 1C) diffraction-limited spots, indicating binding without forming large clusters. The spot sizes and GlyR concentrations (Fig. S1A, as calculated by eGFP fluorescence intensity, see methods for details) did not significantly change with or without GPHN_E (Fig. 1E, F black and blue).

**Figure 1.**
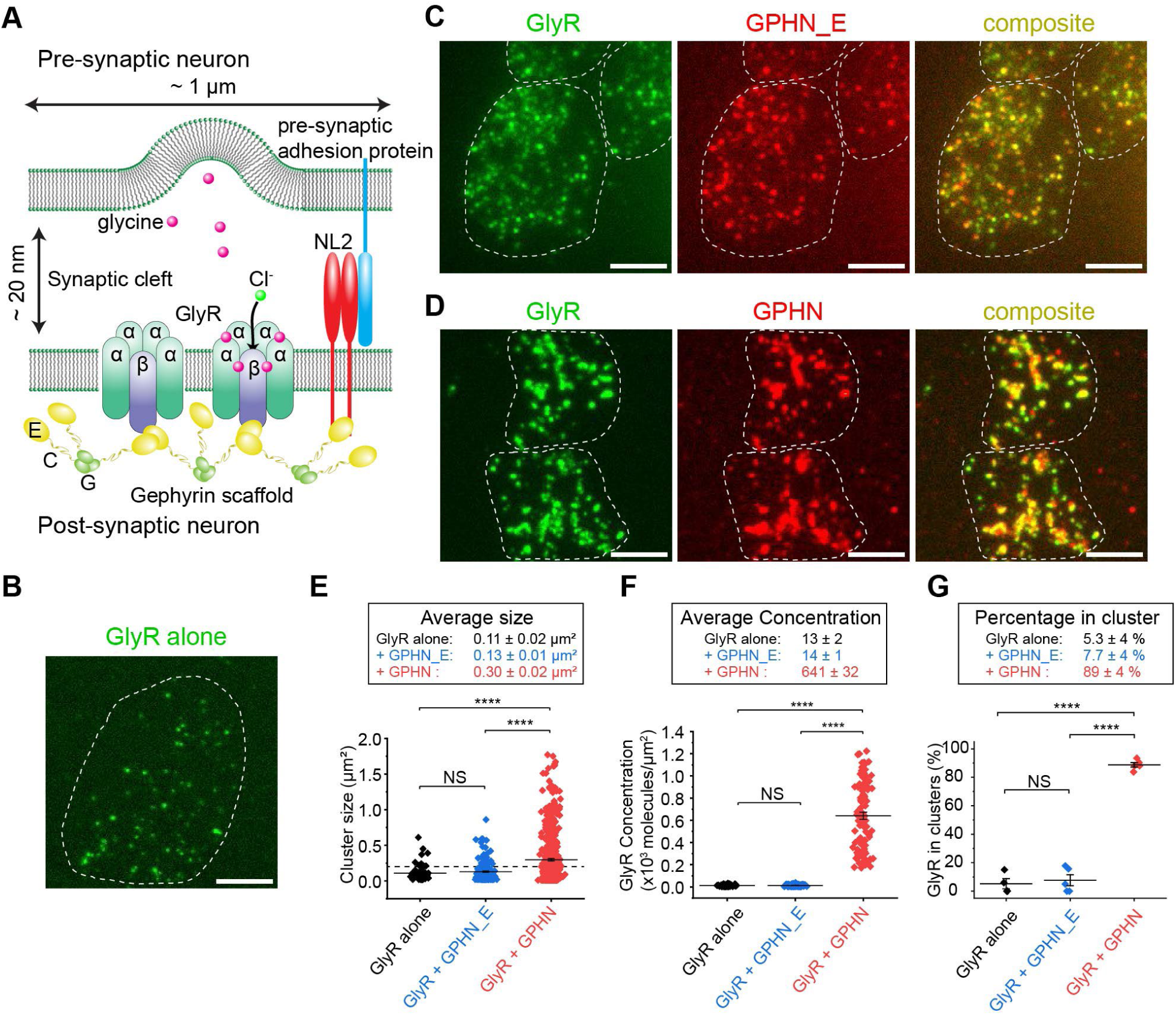
Quantitative analysis of α2β GlyR co-expressed with GPHN_E or GPHN in HEK293 cells. Scale bars: 5 μm **(A)** Illustration of GlyR-gephyrin clusters at the synapse. E, C and G domains of gephyrin are indicated. **(B)** Typical TIRF micrograph of α2β GlyR expressed in HEK293T cells, with single eGFP fused in the β subunit. **(C)** Micrographs of α2β GlyR co-expressed with mCherry- fused GPHN_E. **(D)** Micrographs of α2β GlyR co-expressed with mCherry- fused full-length GPHE. **(E)** Cluster/spot sizes of GlyR alone (black), GlyR + GPHN_E (blue), and GlyR + GPHN (red) expressed in HEK293T cells. Horizontal bars indicate mean ± S.E.M. Dashed line (0.149 µM^2^) shows threshold for defining clustered GlyR. For unpaired t-tests, NS: not significant, **** P<0.0001.) **(F)** Number of GlyRs per µm^2^ of membrane. black: GlyR alone, blue: GlyR + GPHN_E, red: GlyR + GPHN. Horizontal bars indicate mean ± S.E.M. **(G)** Percentage of GlyR in clusters. Horizontal bars indicate mean ± S.E.M.

Full length gephyrin (GPHN) induced clustering of GlyR in cellular membrane (Fig. 1D). Co-expression of GlyR with GPHN resulted in the appearance of large, co-localizing clusters of GlyR and GPHN (Fig. 1D). This is consistent with earlier studies showing GPHN being sufficient and necessary for the clustering of GlyR in native neurons or after heterologous expression^10,41^. GPHN significantly increased cluster sizes, resulting in a wide distribution with ∼ 0.3 µm^2^ (∼ 0.55 µm linear size) average and up to ∼ 1.7 µm^2^ (∼ 1.3 µm linear size) maximum (Fig. 1E red), much beyond diffraction limit (∼ 0.3 µm linear). Coincidentally, the GlyR concentrations also dramatically increased, reaching up to 641 ± 32 GlyRs per µm^2^ (Fig. 1F, please see methods and Fig. S1 for details of quantitative analysis).

GPHN clusters GlyR with high efficiency. Higher-order clustering was defined as fluorescent detections significantly larger than GlyR alone. Since 95% of GlyR alone detections were smaller than 14 pixels (∼0.15 µm^2^, see Fig. S1F), spots with more than 14 pixels were defined as clustered. Under this criterion, ∼ 90% of GlyR is clustered in the presence of GPHN, compared to less than 8% with GPHN_E and ∼ 5% for GlyR alone. Clearly, GlyR clustering by GPHN is recapitulated through heterologous expression in HEK293 cells.

### NL2 further promotes GlyR cluster by GPHN

Co-expression of NL2 increased both GlyR-GPHN cluster sizes and GlyR concentrations. Figure 2A shows two transiently transfected cells that had differential NL2 spatial distribution. The cell on top had more NL2 toward the right side of the micrograph, which co-localized with GlyR-GPHN in visually larger clusters. The bottom cell had more homogeneous NL2 expression and large clusters. The area of clusters averaged ∼ 0.4 µm^2^ and reached ∼ 3.3 µm^2^ in some cases (Fig. 2B). GlyR concentration also increased, averaging at 1414 ± 90 GlyRs / µm^2^ and reaching up to over 4000 GlyRs / µm^2^. This is a significant increase in both size and GlyR concentration compared to GlyR and GPHN alone (Fig. 1). Apparently, NL2 promotes the formation of larger and more concentrated GlyR clusters. On the other hand, the fraction of GlyR in clusters did not change significantly (Fig. 2D).

**Figure 2.**
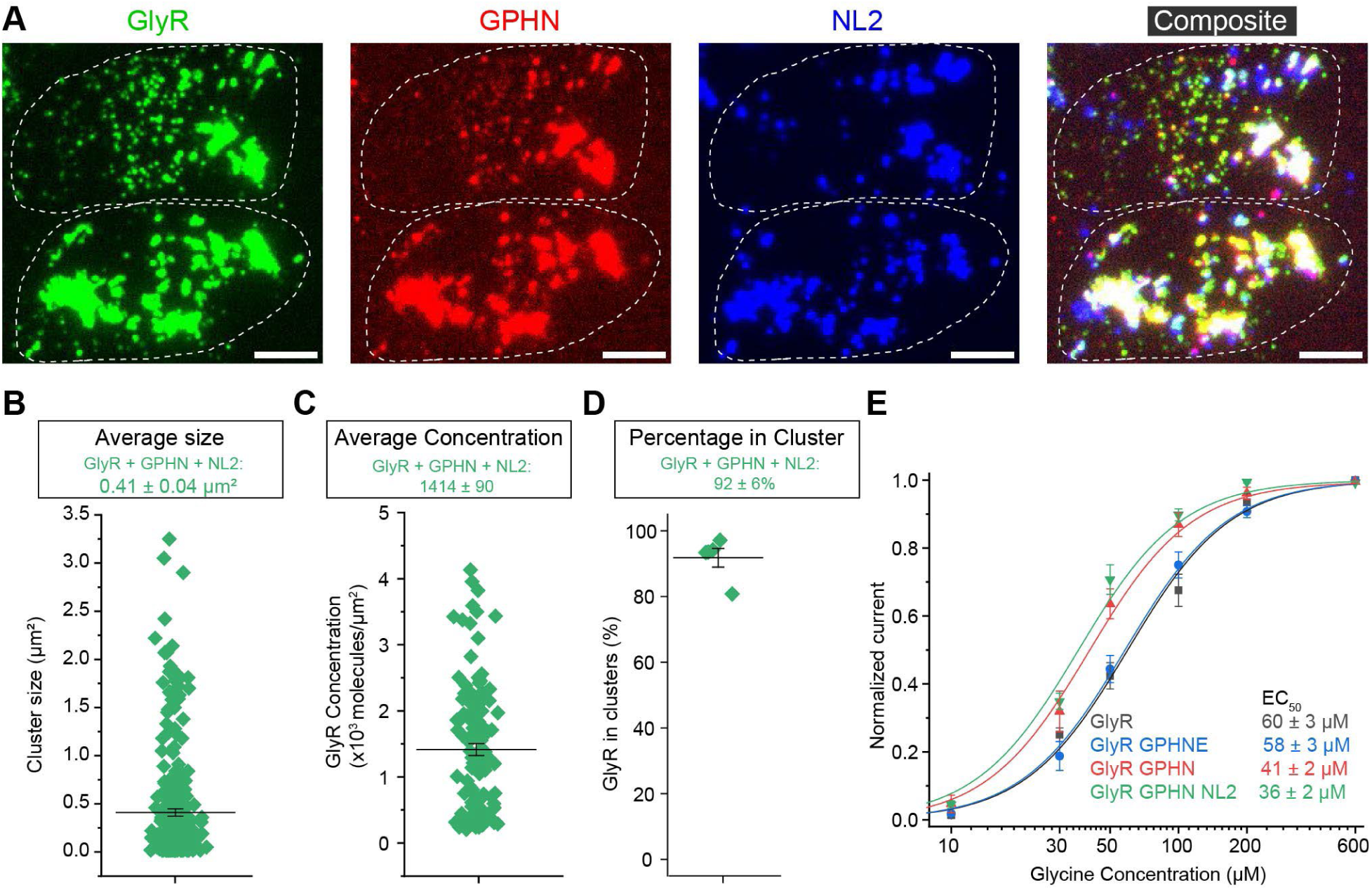
Clustering of GlyR by GPHN increased glycine affinity and is further promoted by NL2. **(A)** Typical TIRF micrographs of α2β GlyR co-expressed with mCherry-GPHN and CF660C labeled NL2. (Scale bar: 5 μm). **(B)** Cluster sizes of GlyR GPHN NL2 co-expressed in HEK293T cells. (B) through (E) horizontal bars indicate mean ± S.E.M. Unpaired t-test P<0.0001 compared to GlyR GPHN **(C)** Numbers of GlyRs per µm^2^. **(D)** Percentage of GlyR in clusters. **(E)** Glycine dose response measured using whole-cell patch clamp in HEK293 cells transfected with GlyR (black), GlyR + GPHNE (blue), GlyR + GPHN (red) or GlyR + GPHN + NL2 (green). (n = 6∼10 cells). Solid lines are Hill fits, with EC50 shown.

Apparent glycine affinity increased as GlyR forms clusters (Fig. 2E and Fig. S2). The glycine EC50 for α2β GlyR when expressed alone is 60 ± 3 µM (Fig. 2E, black). With the co-expression of GPHN_E that binds to GlyR but does not induce clustering (Fig. 1C), EC50 was essentially identical at ∼ 58 ± 3 µM (Fig. 2E blue). Clustering by GPHN (Fig. 1D) reduced EC50 to 41 ± 2 µM (Fig. 2E red), and additional co-expression of NL2 (Fig. 2A) further decreased EC50 to 36 ± 2 µM (Fig. 2E green). Although the change in EC50 is moderate, GlyR activity is much different at lower glycine concentrations due to its sigmoidal dose-response (see Fig. S2 for typical whole-cell recordings, discussed later). Since NL2 increased both cluster sizes and GlyR concentrations, we engineered GPHN to test whether the increase in apparent glycine affinity is resulting from increased cluster size or GlyR concentration.

### GlyR concentration modulates apparent glycine EC50 in clusters

We engineered GPHN for controlling GlyR concentration in clusters. The C domain of GPHN is a semi-flexible linker between the G and E domains, the compactness of which is regulated by multiple phosphorylation sites in vivo^47,48,51^ (Fig. 1A and Fig. 3A). To test how C domain compactness affect GlyR cluster properties in a simplified system, we replaced C domain by flexible linkers of different lengths. These engineered GPHN contained 1 (GE6), 2 (GE12) or 4 (GE24) repeats of SGSGSG peptide sequence. The distance between adjacent GlyRs should change as a function of linker length (Fig 3A). Indeed, engineered GPHN constructs co-localized with GlyR and NL2 upon co-expression, forming large clusters with high fluorescence intensities (Fig. 3B). Quantification of these clusters (Fig. S3) shows a dramatic increase in GlyR concentration (Fig. 3D, from 362 ± 14 to 3710 ± 317 per µm^2^), with a moderate concurrent increase in cluster size (Fig. 3C, from ∼0.2 µm^2^ to ∼0.4 µm^2^).

**Figure 3.**
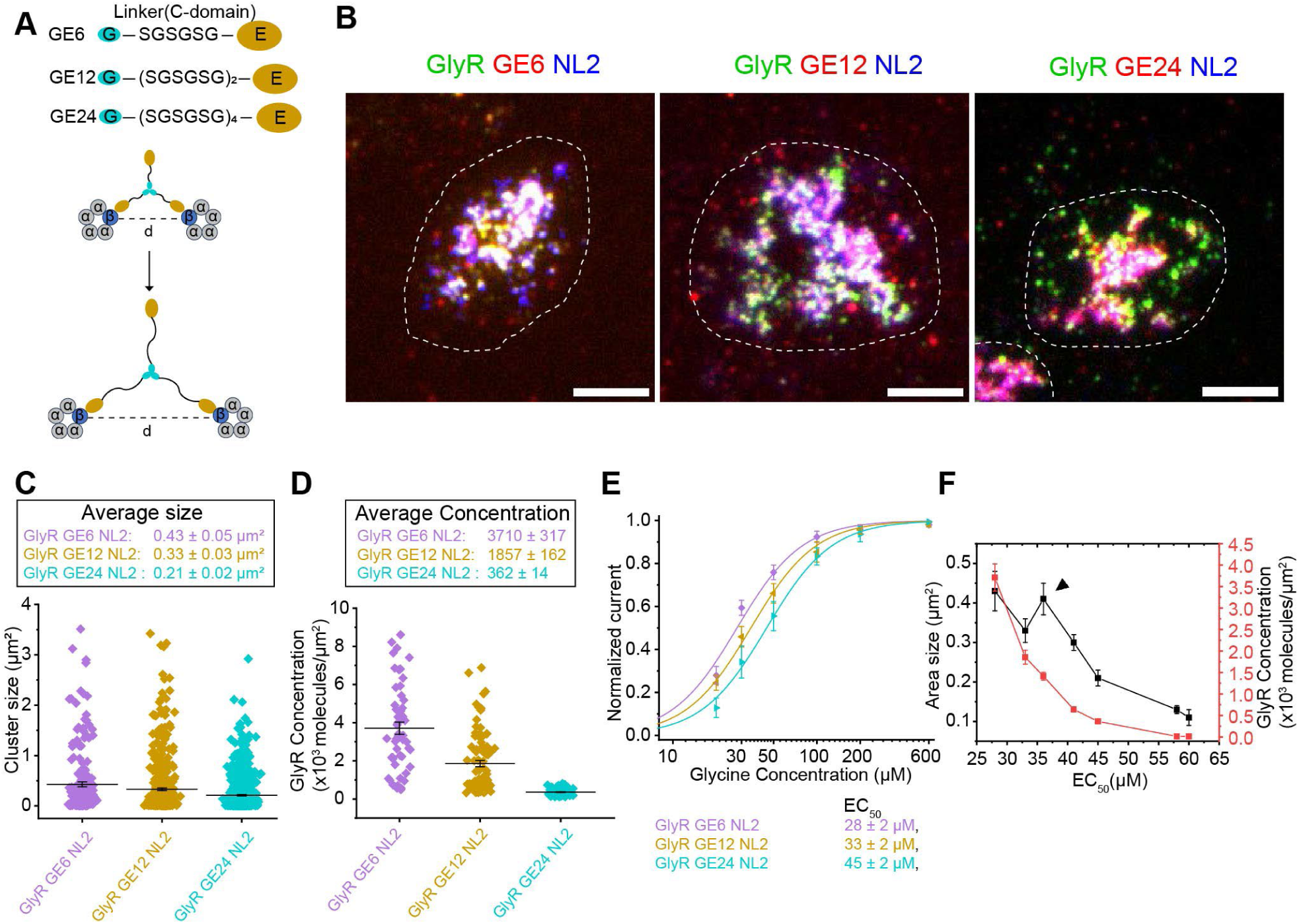
Modulation of GlyR cluster properties and glycine affinity by engineered GPHN. (A) Illustration of GPHN constructs, GE6, GE 12 and GE24, with C-domain replaced with linkers of indicated lengths. Distances (d) between clustered GlyRs are indicated. (B) TIRF micrographs of α2β GlyR (green) co-expressed with GE6/GE12/GE24 (red) and NL2 (blue) shown as merged pseudo-colors. (Scale bar 5 μm). (C) Cluster sizes of GlyR GE6/GE12/GE24 NL2 co-expressed in HEK293T cells. (C) through (F) Horizontal bars indicate mean ± S.E.M. (D) Number of GlyRs per µm^2^ in clusters containing GE6/GE12/GE24 and NL2. (E) Glycine dose response of GlyR clustered with NL2 and GE6 (purple), GE12 (yellow) and GE24 (cyan). (n = 7∼10 cells). Solid lines are Hill fits, with EC50 listed. (F) Plot of cluster size (black) and GlyR concentration (red) against glycine EC50. The black triangle indicated area size fitting outlier.

Higher GlyR concentrations correlated with higher apparent glycine affinities. Glycine EC50 decreased as the linker length decreased in GE24, GE12 and GE6 constructs, with corresponding values of 45 ± 2 µM, 33 ± 2 µM and 28 ± 2 µM (Fig. 3E). Taking together the measurements of GlyR-GPHN (Fig. 1D), or GlyR-GPHN plus NL2 (Fig. 2A), we derived a plot of glycine EC50 against cluster sizes and GlyR concentration (Fig. 3F). Clearly, EC50 increased (affinity decreased) in monotonically as GlyR concentration decreased (Fig. 3F, red). Although higher GlyR concentrations usually correlated with larger cluster sizes, GlyR-GPHN-NL2 clusters had larger size but smaller GlyR concentration compared with GlyR-GE12-NL2 clusters (Fig. 3F, black triangle), correlating with a higher EC50 for GlyR-GPHN-NL2. This suggests that GlyR concentration, instead of cluster size, is the dominant factor for glycine affinity modulation.

### Clustering decreased ligand kinetics through fast re-binding

When GlyR concentrations are sufficiently high, dissociated ligands from one GlyR may quickly re-bind to neighboring GlyRs, slowing down apparent off rates (Fig. 4A). This effect is difficult to measure directly for glycine due to its low affinity and correspondent short dissociation time constant (∼0.1 ms assuming a 10^8^ M^-1^ s^-1^ diffusion-limited on rate^52^ at ∼ 0.1 mM affinity, Fig. 2E). However, strychnine, an orthosteric antagonist that binds with nano-molar affinity^35,53,54^, has much slower off rates, on the order of 10 s, suitable for experimental characterization.

**Figure 4.**
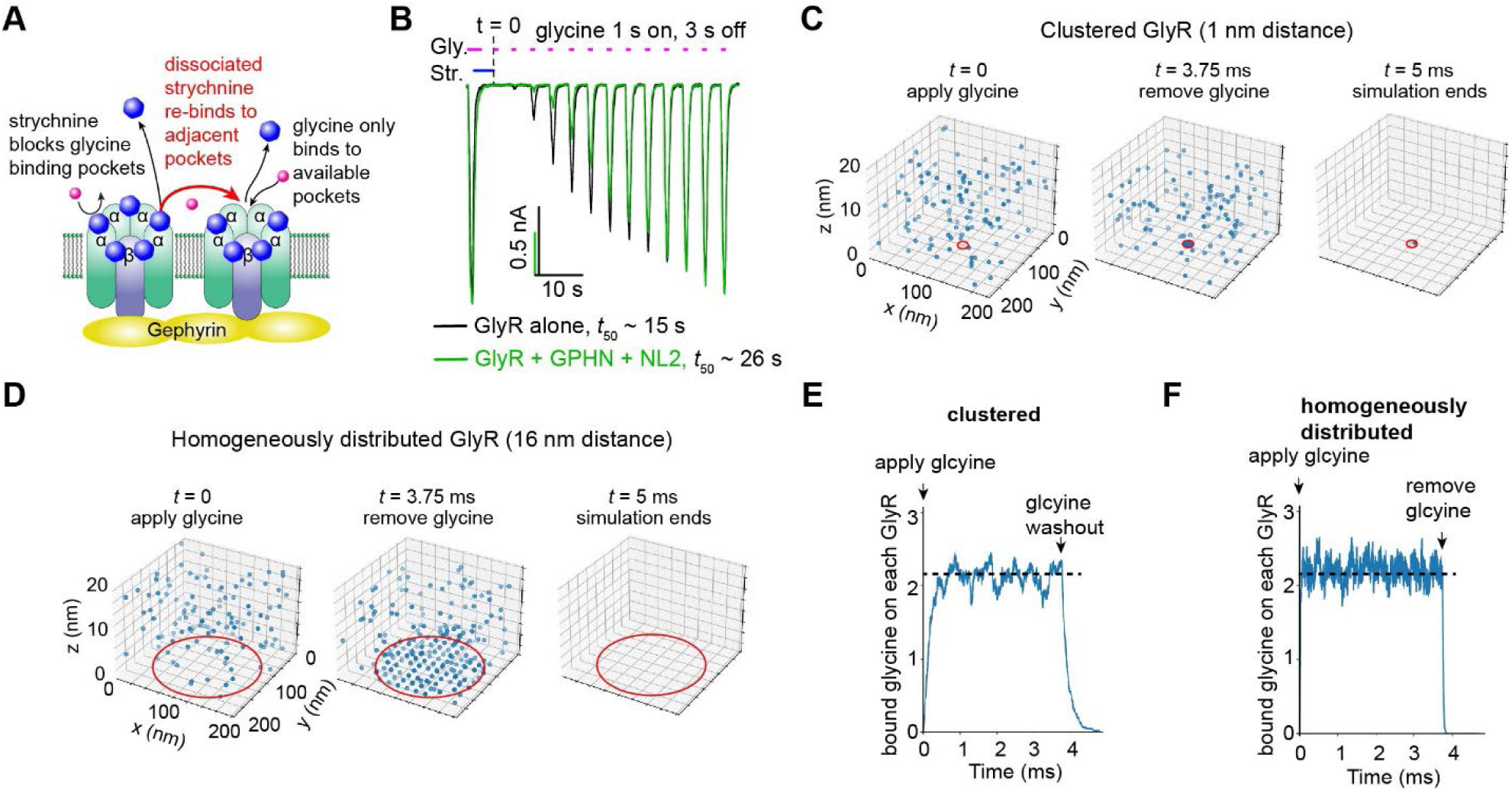
Fast re-binding slows apparent kinetics without shifting equilibrium. **(A)** Illustration of strychnine off-time measurements, highlighting re-binding in red. **(B)** Overlay of typical whole cell recordings of clustered (green) and not clustered (black) GlyR during strychnine off time measurements. Str.: strychnine 10 µM, Gly.: glycine 100μM. Average half-off times, t50, were shown (n = 5-10 cells). **(C)** Snapshots of simulation for glycine binding and dissociation to clustered GlyRs. Each blue dot is a glycine molecule. The red circle shows the region within which GlyRs are placed. **(D)** Snapshots of simulation for glycine binding and dissociation to homogeneously distributed GlyRs. Each blue dot is a glycine molecule. The red circle shows the region within which GlyRs are placed. **(E)** Average number of glycine molecules bound to each GlyR as a function of simulation time, for clustered GlyR. **(F)** Average number of glycine molecules bound to each GlyR as a function of simulation time, for homogeneously distributed GlyR.

Clustering of GlyR by GPHN and NL2 slowed apparent strychnine dissociation (Fig. 4B). 10 µM strychnine (∼ 1, 000 x Kd) was applied at the start of experiment to ensure near complete occupation of orthosteric pockets, as indicated by non-detectable currents in the presence of 100 µM glycine (∼ 2 x EC50). Glycine was applied every 4 seconds to evaluate the dissociation of strychnine (recovery of glycine-inducible current). Without clustering, the half-time, *t50*, for strychnine dissociation was 15 ± 2s, while with clustering, *t50* was 26 ± 1s. Clearly, the concentration of GlyRs in clusters induced by GPHN and NL2 is sufficient for significant re-binding effects to show.

Although the kinetics of glycine binding and dissociation is difficult to experimentally measure, we show similar slowing of kinetics by clustering through simulation (Fig. 4C-F, see methods for details). We evaluated glycine binding to clustered GlyRs (Fig. 4C, 98 GlyRs with 1 nm spacing at center of synaptic membrane, indicated by red circle), or homogeneously distributed GlyRs (Fig. 4D, 16 nm distance, red circle) in a synaptic space measuring 200 nm in width and 20 nm in height. 200 free-diffusing glycine molecules (∼ EC40) were applied at the start of the simulation and kept constant, until at 3.75 ms when free glycine molecules were removed. Clearly, both the binding and dissociation of glycine to clustered GlyR (Fig. 4C, E) was slower than non-clustered (Fig. 4D, F). However, the average numbers of glycine bound to each GlyR (∼2.1) at equilibrium were nearly identical for both cluster and non-clustered case. These calculations explain the slowed binding kinetics in clustered GlyRs through fast re-binding (Fig. 4B), and shows that binding equilibrium does not shift. The measured glycine affinity increase upon clustering (Fig. 2, 3) are near-equilibrium measurements (∼ 0.1 s time scale, compared to ∼0.1 ms glycine binding kinetics)^55^ and thus unlikely resulting from re- binding effects.

### Electrostatics underly clustering-induced glycine affinity modulation

A stretch of positively charged amino acid residues is found at the N-terminus of GlyR β subunit (Fig. 5A). This sequence (aa 1-13) contains 8 lysine residues and a single glutamate residue, contributing to a net total charge of around +7 on the extracellular side. Cl^-^ is negatively charged at physiological pH. Glycine has an isoelectric point of ∼ pH 6.0, and is therefore slightly negatively charged, especially at pHs above 7.5. Local concentration of these anions by the positive charges of GlyR may contribute to elevated apparent glycine affinity. To test this, we evaluated glycine affinities after removal, or reversal of these charges.

**Figure 5.**
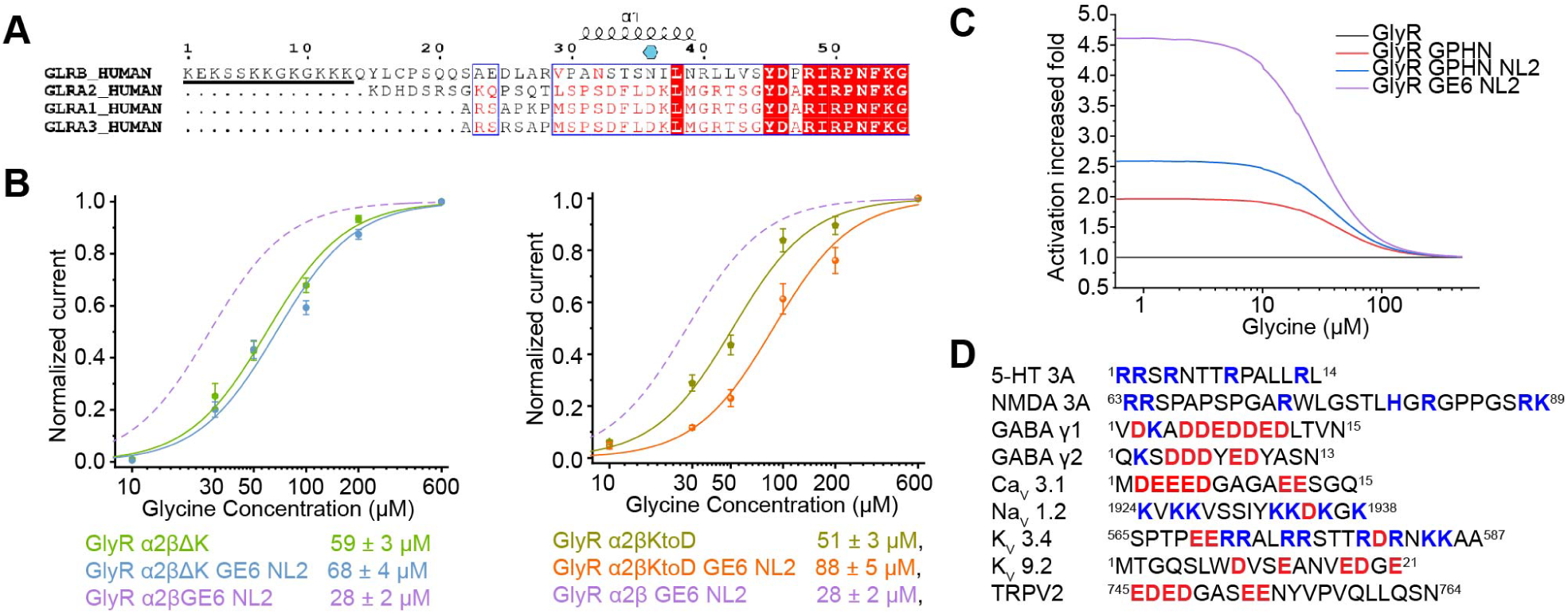
Electrostatic charges underly ion channel function modulation by clustering (A) Sequence alignment among GlyR subunit types show β subunit contains N-terminal sequence with multiple positively charged residues (underlined). Residue conservation was indicated by blue rectangle and red highlights. Secondary structure elements were indicated by helices for α-helices above the alignment. N-linked glycosylation site of GlyR β was indicated by cyan hexagon. Protein sequences aligned: human GLRB (Uniprot P48167), human GLRA2 (Uniprot P23416), human GLRA1 (Uniprot P23415), human GLRA3 (Uniprot O75311). (B) Glycine dose response of α2βΔK GlyR (left) and α2βKtoD GlyR (right). Data points with S.E.M. (n = 6∼8 cells) were plotted with Hill fits, and EC50 listed below. (C) Fold increase of GlyR activity upon clustering as a function of glycine concentrations. (D) Examples of ion channels containing charged sequences near the N- or C-terminus: 5-HT3A(Uniprot: P46098), glutamate receptor NMDA 3A (Uniprot: Q8TCU5) GABAA γ1 (Uniprot: Q8N1C3), GABAA γ2 (Uniprot: P18507), CaV3.1(Uniprot: O43497),NaV1.2 (Uniprot: Q99250), KV3.4 (Uniprot: Q03721), KV9.2 (Uniprot: Q9ULS6), TRPV2 (Uniprot:). Positively- and negatively- charged amino acids are highlighted in blue and red, respectively.

Removal of the N-terminal charges of the GlyR β subunit abolished functional effects of clustering. Without clustering, the α2β GlyR with removed β:aa1-13 (α2βΔK) had a similar glycine EC50 = 59 ± 3 µM (Fig. 5B left, green) as wild type (EC50 = 60 ± 3 µM, Fig. 2E black). Co-expression of α2βΔK with GPHN with the shortest linker (GE6, Fig. 3A) and NL2 induced clustering (Fig. S4A) but did not increase glycine affinity (EC50 = 68 ± 4 µM, Fig. 5B left, blue). This contrasts the wild type where clustering decreased EC50 to ∼ 38 µM (Fig. 3E and Fig. 5B left, purple).

Reversal of the GlyR β subunit N-terminal charges reversed functional effects. We generated a β subunit with all lysine residues in aa1-13 mutated to aspartate residues (βKtoD), effectively reversing the charges. The α2βKtoD GlyR had similar glycine EC50 = 51 ± 3 µM before clustering. Although it clustered with GE6 and NL2 like wild-type (Fig. S4B), the glycine EC50 increased upon clustering, to a value of 88 ± 5 µM (Fig. 5B, right). This is opposite to wild-type, where glycine EC50 decreased from ∼ 60 µM to ∼ 30 µM (Fig. 5B right, purple, Fig. 2E and Fig. 3E).

The seemingly small change in EC50 (∼ 2 folds) may lead to large GlyR activity differences especially at lower glycine concentrations (Fig. 5C). This is due to the cooperative nature of glycine activation, with Hill slopes around 2^55–57^. GlyR activity is very low at glycine concentrations lower than the rising phase of the sigmoidal dose-response curve, meaning a large fold-change with moderate EC50 shift (Fig. 5C). Indeed, GlyR activity may be increased by nearly 5 folds at glycine concentrations below 10 µM (Fig. 5C purple). These differences decrease as glycine concentrations increase and becomes negligible at concentrations much above EC50.

Charged stretches of amino acids are commonly found in ion channels (Fig. 5D). Examples include neurotransmitter receptors 5-HT3A, NMDA3A and GABAA γ1/2, voltage gated sodium channel Nav1.2, potassium channels Kv3.4 and Kv9.2, calcium channel Cav3.1 and transient receptor potential channel TRPV2. Ion channels are naturally modulated by local electrostatics since they conduct charged ions. In addition, many ligands of ion channels are charged at physiological pH. It is tempting to speculate that enhancement of local electrostatics through high concentration clustering may constitute a universal mechanism of ion channel function modulation (Fig. 6).

**Figure 6.**
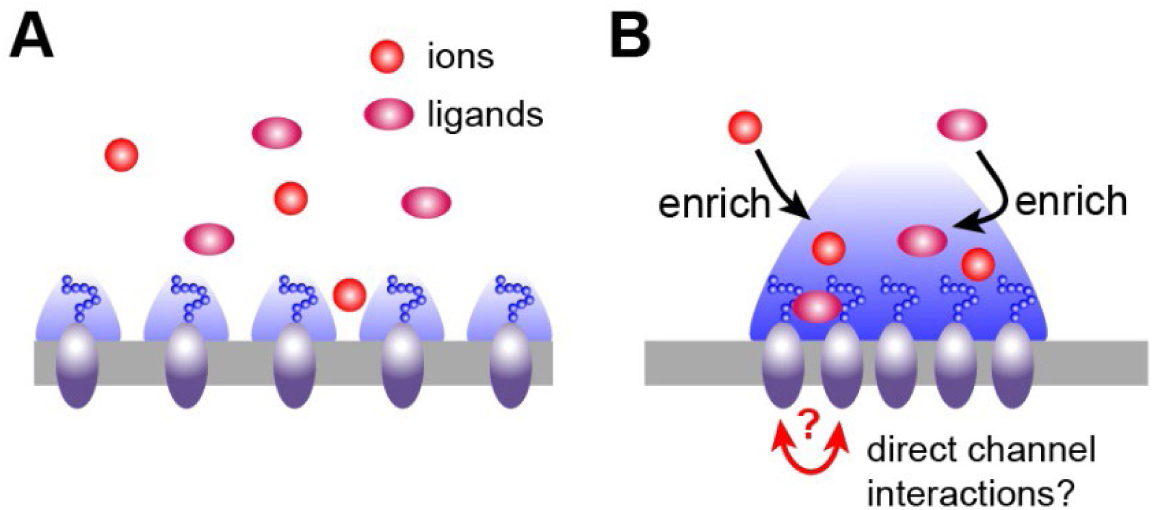
Illustration of electrostatically driven local enrichment of ions and ligands upon GlyR clustering. (A) Without clustering, electrostatics fields from individual GlyRs are weaker and smaller. (B) Once clustered, local charge density increases, increasing both intensity and dimension of relevant electrostatic fields, locally enriching countercharges including ions and charged ligands.

## Discussion

We showed that co-expression of GlyR and GPHN results in micron-sized clusters in HEK293 cell membrane, the size and GlyR concentration of which is further increased by the trans-synaptic adhesion protein NL2. Clustering of GlyR increased apparent glycine affinity monotonically with the increase of GlyR concentration. We further showed that fast re-binding in the clusters slowed kinetics but did not change binding equilibrium and thus unlikely responsible for increased glycine affinity. Instead, we identified a positively-charge N-terminus loop on the β subunit of GlyR being essential for clustering-induced glycine affinity increase. Since charged sequences are commonly found in multiple types of ion channels, clustering-induced change of local electrostatics may be a universal mechanism for ion channel function modulation.

High-concentration clustering of charge-carrying GlyR and other ion channels leads to locally enhanced electrostatic fields (Fig. 6). The electrostatic filed intensity can be simply conceptualized by a charged disc model, with field strength on the symmetrical axis 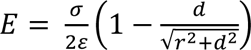, where *σ* is charge density (q/area), *ε* is permittivity, *d* is distance from disc and *r* is disc radius. Without clustering, the charge density is low, resulting in limited electrostatic fields (Fig. 6A). Clustering of GlyR increases charge density, resulting in a much higher local electrostatic field strength (Fig. 6B). This stronger field in turn attracts counterions and locally enriches charged species including Cl^-^ and glycine, shifting equilibrium of ligand binding and changes apparent affinity.

The GlyR β subunit carrying the N-terminal positively charged amino-acid residues (Fig. 5A) coincides with its essential role for GlyR clustering^42,58^ – only the β subunit contains the GPHN-binding site. The functional roles of the N-terminal charges have not been identified before, and we show here changes apparent glycine affinity upon GlyR clustering. In cells, the compactness of GPHN is regulated by phosphorylation^46–49,51^, suggesting that modulating GlyR function through post-translational modification of GPHN is an unidentified mechanism of regulation uniquely enabled by the GlyR β subunit.

GlyR clusters have properties reminiscent of liquid-liquid phase separated systems ^59,60^. In addition, several lines of evidences supports phase separation being an important mechanism in organizing ion channels at the synapse in a near 2D setting delimited on the membrane^59–66^. Biomolecular condensates change the concentration and kinetics (dwell times) of molecules to elicit functional effects that fulfil physiological function^62,67,68^. Since there is only one β subunit in each GlyR^35,36^, multivalent GPHN/NL2 constitute the backbone/scaffold that dictates clustering^35,61^. The function, namely ion conduction, is purely through the GlyRs that are activated through the binding of glycine and regulated by Cl^-^ concentration. Without additional mechanisms that increase free glycine and/or Cl^-^ concentrations, clustering and accompanied fast re-binding effect^69,70^ slows ligand-binding kinetics but does not shift chemical equilibrium. However, concentration of immobilized charges (on channels) dramatically increases surface charge density and attract counterions to reach equilibrium. This provides a mechanism that locally enriches ions and charged small ligands, therefore modulating equilibrium ion channel function in high density clustered likely formed through liquid-liquid phase separation.

Whether adjacent GlyRs directly interact with each other is unclear. The clustering concentrates GlyR up to 8000 per µm^2^. This corresponds to an average distance of 11 nm between centers of adjacent GlyRs. Considering a GlyR diameter of ∼ 10 nm, collisions resulting from diffusion should happen frequently at these concentrations. Further structural characterization of the GlyR-GPHN clusters is required to understand possible GlyR-GlyR interactions, as well as specific spatial organizations that may promote functional modulation or structural stability.

## Acknowledgments

We thank Robbie Boyed for preparation of cell cultures, Dr. Michael Rosen for helpful discussions, and members of the Wang laboratory for technical assistance. Cryo-EM data was collected at the University of Texas Southwestern Medical Center Cryo-EM Facility, which is funded by the CPRIT Core Facility Support Award RP170644. This work is supported by NIH grant 1R35GM146860 and the McKnight Scholar Award to W.W.

## Experimental model and subject details

### Cell lines

HEK293T cells (ATCC CRL-3216) used for TIRF imaging and electrophysiology were cultured in Freestyle 293 expression medium (GIBCO) supplemented with 1% FBS at 37°C in 5% CO2 incubator.

### Method details

#### Molecular Biology and constructs

The human GlyR α2 or β gene was cloned into a modified plasmid vector pBM ^71^, as detailed described previously^35^. Briefly, the M3/M4 loop (Q317-K381) of GlyR α2 subunit was substituted by GSSG peptide. The GlyR β construct contained the following modifications: region encoding M3/M4 loop (residues N334-N377) was removed and an monomeric enhanced green fluorescent protein (mEGFP)^72^ was inserted. These constructs are used throughout this work and denoted α2 and β for simplicity unless otherwise specified. The GlyR β N-terminal poly-lysine mutants were cloned based on the previous construct. The N-terminal poly-lysine was deleted (denoted as βΔK) or mutated to Asp (denoted as βKtoD). Human GPHN (NM_020806.4) full-length or E domain (M351-L769) was cloned into the pBM vector with an N-terminal mCherry and PreScission protease site (LEVLFQ/GP)^73^. Based on the GPHN full-length construct, the linker (E182-R350) between G domain and E domain was deleted and substituted by SGSGSG, (SGSGSG)2 or (SGSGSG)4 flexible linker. These constructs were denoted as GE6, GE12 or GE24 based on the flexible liner residues numbers. The rat NLGN2 (NM_053992) with a HA-tag (YPYDVPDYA) was kindly provided by Alice Y. Ting (Stanford University)^74^ and cloned into pBM vector. Based on the NLGN2 construct, a Halo-tag^75^ was inserted at the C-terminal of NLGN2 for electrophysiology study(the construct denoted as NL2- Halo). Gibson Assembly was used to make these constructs.

### TIRF imaging

Coverslips were coated by 0.1mg/mL collagen. Plasmids bearing α2:β at 1:3 ratio (α2:β:GPHN at 1:3:1 ratio) were transfected into HEK293T cells following manufacturer’s protocol by lipofectamine 3000 (Invitrogen) for expression GlyRs on coverslips. The transfected cells were kept in CO2 incubator at 37°C for overnight expression. Cells were unroofed^76^ and fixed prior to imaging as follows: After 3 times wash at room temperature (RT) with Phosphate-Buffered Saline PBS (10 mM Na2HPO4, 1.8 mM NaH2PO4, 137 mM NaCl, 3 mM KCl), cells were sonicated at 20% power for 1 s using a Branson SFX550 sonicator equipped with a flat microtip in ice-cold PBS supplemented with 2% para-formaldehyde (PFA), 2 mM MgCl2 and 1 mM CaCl2. Unroofed cells were washed quickly with ice-cold PBS and immediately switched into PBS containing 4% PFA at RT and fixed for 15min. After 3 times washing in PBS for 10 min each, the cells were imaged within 2 hours. Total internal reflection illumination was achieved using an in-house built prism-type system using the Gem 488/561 laser (Laser Quantum) and components from Thorlabs at 50W. A 1.5 OD neutral density filter was used during searching of cells to minimize photobleaching. A Leica DM6 FS microscope equipped with a 63x 1.2 NA CS2 water immersion objective and a Hamamatsu flash 4.0 V3 camera was used for imaging.

Movies were collected using a PC running Metamorph (Molecular Devices) with 100 ms exposure for each frame and total exposure of 50 s.

### Immunofluorescence Staining for TIRF imaging

Plasmids bearing GlyR α2, β, GPHN, NLGN2 were transiently transfected into HEK293T cells by Lipofectamine 3000 (Invitrogen) at α2:β:GPHN:NLGN2 DNA 1:3:1:1 ratio. After transfection, cells were cultured at 37°C for overnight culture. The cells were unroofed and fixed as previously described. Then the cells were blocked with blocking buffer (PBS + 1% BSA).

Primary antibody (HA-Tag C29F4 Rabbit mAb) was diluted with fresh blocking buffer at 1:2000 ratio. The cells on the coverslip were incubated overnight at 4°C. Then the cells were rinsed four times by PBS. The secondary antibody (Goat Anti-Rabbit IgG, Highly Cross-Adsorbed, labelled with CF660C dyes, Biotium, Cat # 20369) was diluted with blocking buffer to final concentration of 1 ug/mL. The cells were incubated with secondary antibody solution for 1 hour at room temperature (protected from light). Then the coverslip was rinsed by PBS buffer several times before TIRF imaging. The Total internal reflection illumination was achieved using an in-house built prism-type system using the Gem 488/561/660 laser. The other microscopy settings were the same as previously described.

### Images analysis

The regions of interest were cropped using ImageJ^77^ and analyzed. GlyR GFP intensities were detected by LoG (Laplacian of Gaussian) filter for micrographs of GlyR, and GlyR + GPHN_E samples using TrackMate ^78^. Intensities were plotted as histogram and fitted by bimodal gaussian equation 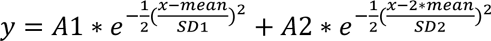, where mean is the first peak center, SD1 and SD2 are the variances. To analysis the GlyR size, firstly, the cropped images were processed with Bernsen’s ^79^ or Phansalkar’s ^80^ local thresholding method (Figure S1). Secondly, the number of pixels and average intensities of each ROI (GFP channel) were analyzed and saved. The background intensity was also measured for each micrograph. For later intensity analysis, the background intensity was deducted. Thirdly, the statistics of area at different conditions (GlyR, GlyR GPHNE, GlyR GPHN etc.) were calculated by OriginPro (OriginLab). These statistics included mean of area, S.E.M. of area, lower confidence interval (C.I.) (95%), upper confidence interval (95%). Depending on these data, ROIs larger than 20 pixels (0.2μm^2^, each pixel distance is 0.103μm) were defined as clusters. Finally, the percentage of GlyR in cluster was calculated. To calculate the number of GlyR molecules per μm^2^, the intensities, *I,* and pixel number, n, of each ROI are listed. Total number of GlyRs in each ROI was calculated by *N* = *I* / *I0*, where *I0* is the single GFP fluorescence calculated in Figure S1. Glycine concentration is then calculated by 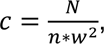, where *w* is the pixel size (*w* = 0.103 μm for our imaging system). Cluster sizes and GlyR concentrations are plotted using OriginPro (OriginLab).

### Whole cell patch clamp

Plasmids bearing GlyR GPHN and NL2 constructs were transiently transfected into HEK293T cells by Lipofectamine 3000 (Invitrogen) following manufacturer’s protocol at the same ratio as previously described. For each 35 mm dish, 1.5 μg of DNA was transfected. After transfection, cells were cultured at 37°C overnight culture. Whole-cell recordings were performed at room-temperature. The bath solution contained (in mM): 10 HEPES pH 7.4, 10 KCl, 125 NaCl, 2 MgCl2, 1 CaCl2 and 10 glucose. The pipette solution contained (in mM): 10 HEPES pH 7.4, 150 KCl, 5 NaCl, 2 MgCl2, 1 CaCl2 and 5 EGTA. Borosilicate glass pipettes with resistance between 3∼8 MΩ were used. A Digidata 1550B digitizer (Molecular Devices) was connected to an Axopatch 200B amplifier (Molecular Devices) for data acquisition. Analog signals were filtered at 1 kHz and subsequently sampled at 20 kHz and stored on a computer running pClamp 10.5 software. Membrane was held at −80 mV during perfusion of ligands to record GlyR current. For the GlyR recordings, as the GlyR β construct had an EGFP insertion, we used GFP fluorescence to identify the cells expressing the GlyR β subunit. For the GlyR GPHN recordings, as the GlyR β construct had an EGFP insertion and the GPHN had an N-terminal mCherry, we used GFP and mCherry fluorescence to identify the cells both expressing the GlyR and GPHN. To identify the cells expressing NL2, the NL2-Halo construct was used for co-transfection. Before whole cell patch clamp, fresh medium with 20nM Janelia Fluor 646(JF646) HaloTag Ligand was changed and incubated for 1 hour to label the living cells.

Then we used GFP/mCherry/JF646 fluorescence to identify the cells expressing the GlyR, GPHN and NL2. Hill equation was used to fit the dose-response data and derive the EC50 (*k*) and Hill coefficient (*n*). For glycine activation, we used 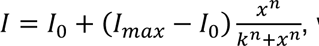, where *I* is current, *I0* is the basal current (spontaneous opening current and leak, very close to 0), *Imax* is the maximum current and *x* is glycine concentration. Data was fitted with software OriginPro (OriginLab, Northampton, MA). Measurements were from 6–10 cells, average and S.E.M. values were calculated for each data point. To measure the strychnine off rate, after breaking into whole cell mode, 100 μM glycine was applied for 2 seconds, followed by addition of 10 μM strychnine for 2 s on top of glycine when GlyR activities were completely inhibited. Strychnine was applied for another additional 2 s in the absence of glycine to ensure complete binding. Afterwards, 1 s of 100 μM glycine and 3 s of bath solution were alternatively applied for 15 rounds till the GlyR current recovered to the level before the application of strychnine. Recordings were analyzed and plotted using pClamp (Molecular Devices).

### Simulation of glycine binding to GlyR at synapse

Simulations were based on simple random walk of glycine in a rectangular simulation box of 200 nm width (synaptic membrane area) and 20 nm height (synaptic cleft). 98 GlyRs were placed at the bottom of the box (post-synaptic membrane) with 1 nm spacing to represent tightly clustered case, or with 16 nm spacing to represent homogenously distributed and non-clustered situation. GlyRs were immobile, and a diffusion constant of 10^3^ µm^2^ s^-1^ was assumed for glycine with a reflective diffusion boundary condition. The on-rate is assumed purely diffusion limited, meaning once glycine diffuse to the position of GlyR, binding happens. The dissociation was assumed a pure Poisson process. 200 glycine molecules, corresponding to ∼ EC40 of GlyR, was randomly placed within the simulation box at the start of simulation, and free glycine molecules kept constant afterward through replenishing at the presynaptic membrane. At 3.75 ms, all free glycine molecules were removed, and any glycine diffused to the presynaptic membrane was removed. The total simulation time was 5 ms. The simulation was performed using a in-house python script written using PyCharm.

