## Supplemental data for "Modulation of heteromeric glycine receptor function through high concentration clustering"

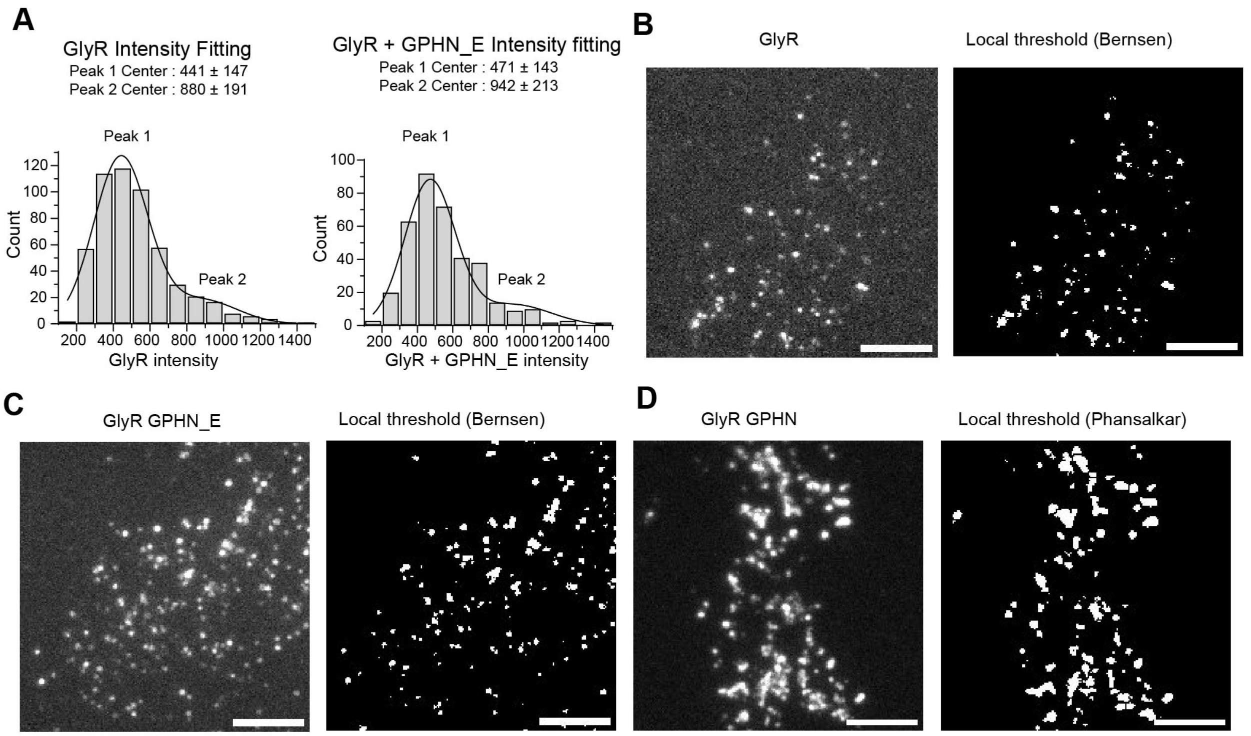


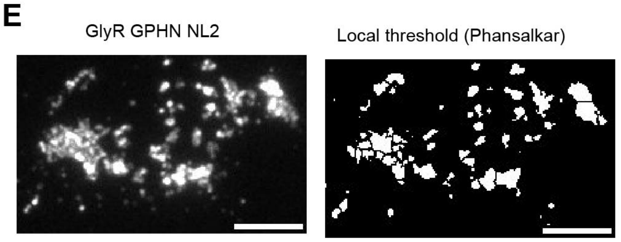


F

| | α2β | α2β GPHN_E | α2β GPHN | α2β GPHN NL2 |
|---|---|---|---|---|
| Mean of area | 11 | 13 | 30 | 41 |
| S.E.M. | 2 | 1 | 2 | 4 |
| Lower C.I., 95% | 8 | 11 | 26 | 33 |
| Upper C.I., 95% | 14 | 15 | 33 | 48 |

**Figure S1. Images processing and quantitative statistical analysis of GlyR co-expressed without/with GPHN_E/GPHN and NL2, related to Figure 1 and Figure 2.**

**(A)** GlyR GFP intensity histogram (bars) and bimodal gaussian fitting (solid lines) of GlyR alone or expressed with GPHN_E.

**(B)** Images of GlyR α2β heteromer and ROIs after Bernsen's local thresholding. (Scale bar 5 μm)

**(C-E)** Images of GlyR and ROIs after Bernsen's or Phansalkar's local thresholding. (C) GlyR GPHN_E co-expression, (D) GlyR GPHN co-expression and (E) GlyR GPHN NL2 co-expression.

**(F)** Area analysis (in pixels) by OriginPro. S.E.M. : standard error of mean, C.I.: confidence interval.

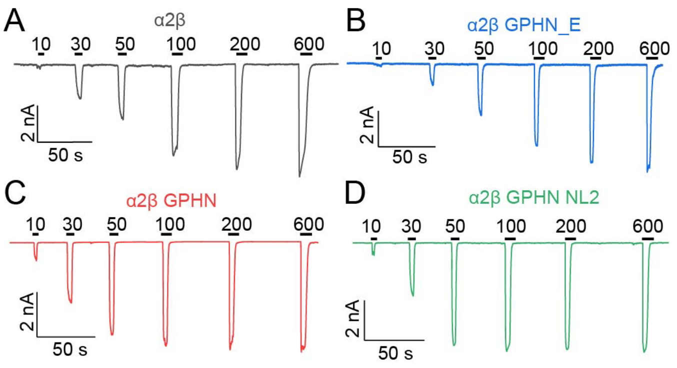


**Figure S2. Whole cell patch clamp electrophysiology of GlyRs at different conditions, related to Figure 2.**

**(A-D)** Representative voltage-clamp recordings of glycine dose response in HEK293 cells transfected with (A) GlyR α2β(Gray), (B) GlyR α2β GPHN_E (blue), (C) GlyR α2β GPHN (red) (D) GlyR α2β GPHN NL2 (green) plasmids.

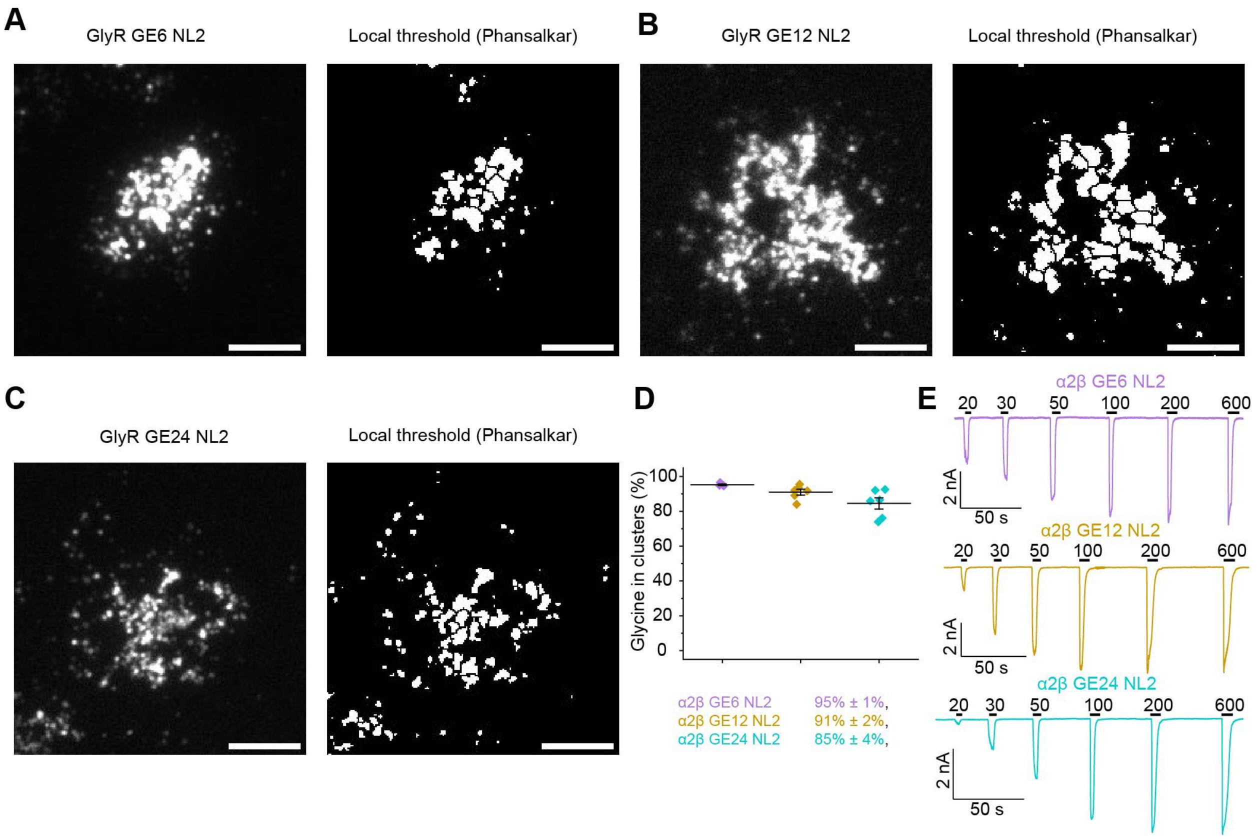


**Figure S3. Images processing and quantitative statistical analysis of GlyR co-expressed with GE6/12/24 and NL2, related to Figure 3.**

**(A-C)** Images of GlyR and ROIs after Phansalkar's local thresholding. (A) GlyR GE6 NL2 co-expression, (B) GlyR GE12 NL2 co-expression and (C) GlyR GE24 NL2 co-expression. Scale bar: 5µm.

**(D)** Percentage of GlyR in clusters when GlyR GE6/12/24 NL2 co-expressed in HEK293T cells. Each symbol represents one cell; horizontal bars and error bars indicate mean and standard error of mean (S.E.M.), respectively.

**(E)** Representative voltage-clamp recordings of glycine dose response in HEK293 cells transfected with GlyR α2β GE6 NL2 (magenta), GlyR α2β GE12 NL2 (brown) and GlyR α2β GE24 NL2 (cyan) plasmids.

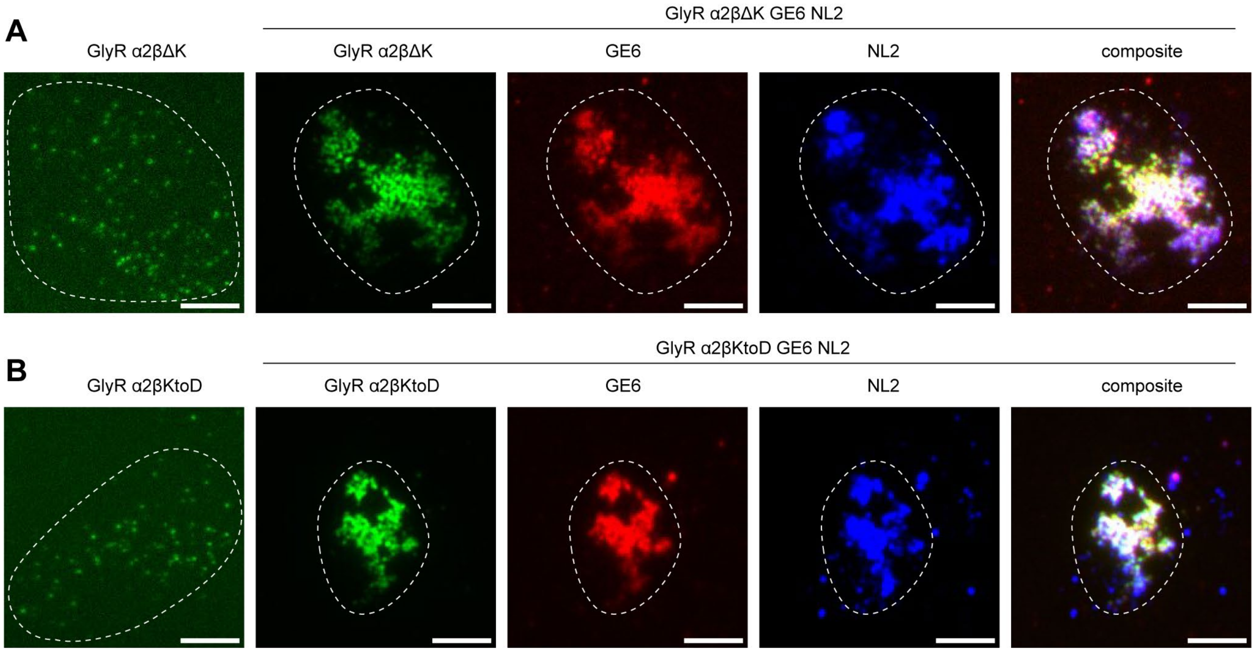


**Figure S4. TIRF imaging of GLRB N-terminal mutation constructs, related to Figure 4.**

**(A)** TIRF imaging of GlyR α2βΔK (left panel) or co-expressed with GE6 and NL2 (right panel) in HEK293T cells. (Scale bar 5 μm).

**(B)** TIRF imaging of GlyR α2βKtoD (left panel) or co-expressed with GE6 and NL2 (right panel) in HEK293T cells. (Scale bar 5 μm).
